# Immune Loss as a Driver of Coexistence During Host-Phage Coevolution

**DOI:** 10.1101/105908

**Authors:** Jake L Weissman, Rayshawn Holmes, Rodolphe Barrangou, Sylvain Moineau, William F Fagan, Bruce Levin, Philip L F Johnson

## Abstract

Bacteria and their viral pathogens face constant pressure for augmented immune and infective capabilities, respectively. Under this reciprocally imposed selective regime, we expect to see a runaway evolutionary arms race, ultimately leading to the extinction of one species. Despite this prediction, in many systems host and pathogen coexist with minimal coevolution even when well-mixed. Previous work explained this puzzling phenomenon by invoking fitness tradeoffs, which can diminish an arms race dynamic. Here we propose that the regular loss of immunity by the bacterial host can also produce host-phage coexistence. We pair a general model of immunity with an experimental and theoretical case study of the CRISPR-Cas immune system to contrast the behavior of tradeoff and loss mechanisms in well-mixed systems. We find that, while both mechanisms can produce stable coexistence, only immune loss does so robustly within realistic parameter ranges.

## 1. Introduction

While the abundance of bacteria observed globally is impressive (Hug *et al.*, 2016; Schloss *et al.*, 2016; Whitman *et al.*, 1998), any apparent microbial dominance is rivaled by the ubiquity, diversity, and abundance of predatory bacteriophages (or “phages”), which target these microbes (Suttle, 2005; Weitz and Wilhelm, 2012; Wigington *et al.*, 2016; Wilhelm and Suttle, 1999; Wommack and Colwell, 2000). As one might expect, phages are powerful modulators of microbial population and evolutionary dynamics, and of the global nutrient cycles these microbes control (Bergh *et al.*, 1989; Bratbak *et al.*, 1990, 1994; Proctor and Fuhrman, 1990; Sieburth *et al.*, 1988; Suttle, 2005; Weinbauer and Rassoulzadegan, 2004; Weitz and Wilhelm, 2012; Whitman *et al.*, 1998; Wilhelm and Suttle, 1999). Despite this ecological importance, we still lack a comprehensive understanding of the dynamical behavior of phage populations. More specifically, it is an open question what processes sustain phages in the long term across habitats.

Bacteria can evade phages using both passive forms of resistance (e.g. receptor loss, modification, and masking) and active immune systems that degrade phages (e.g. restriction-modification systems, CRISPR-Cas) (Labrie *et al.*, 2010). These defenses can incite an escalating arms race dynamic in which host and pathogen each drive the evolution of the other (Rodin and Ratner, 1983a,b). However, basic theory predicts that such an unrestricted arms race will generally be unstable and sensitive to initial conditions (Schrag and Mittler, 1996). Additionally, if phages have limited access to novel escape mutations, an arms race cannot continue indefinitely (Hall *et al.*, 2011; Lenski, 1984; Lenski and Levin, 1985). This leads to an expectation that phage populations will go extinct in the face of host defenses (Lenski and Levin, 1985).

While typically this expectation holds (e.g. van Houte *et al.*, 2016), phages sometimes coexist with their hosts, both in natural (e.g. Gómez and Buckling, 2011; Waterbury and Valois, 1993) and laboratory settings (e.g. Bohannan and Lenski, 1999; Chao *et al.*, 1977; Horne, 1970; Lenski and Levin, 1985; Levin and Udovic, 1977; Paez-Espino *et al.*, 2015; Schrag and Mittler, 1996; Wei *et al.*, 2011). These examples motivate a search for mechanisms to explain the deescalation and eventual cessation of a coevolutionary arms race dynamic, even in the absence of any spatial structure to the environment. Previous authors have identified (1) fluctuating selection and (2) costs of defense as potential drivers of coexistence in well-mixed systems. Here we propose (3) the loss of immunity, wherein the host defense mechanism ceases to function, as an additional mechanism. We focus on intracellular *immunity* (e.g., CRISPR-Cas) in which immune host act as a sink for phages rather than extracellular *resistance* (e.g., receptor modifications), since the former poses more of an obstacle for phages and thus more of a puzzle for explaining long-term coexistence.

Under a fluctuating selection dynamic, frequencies of immune and infective alleles in the respective host and phage populations cycle over time (Agrawal and Lively, 2002; Gandon *et al.*, 2008; Van Valen, 1973, 1974). That is, old, rare genotypes periodically reemerge because the dominant host or pathogen genotype faces negative frequency dependent selection. Fluctuating selection is likely in situations where host immune and phage infectivity phenotypes match up in a one-to-one “lock and key” type manner (Agrawal and Lively, 2002), and there is evidence that arms races do give way to fluctuating selection in some host-phage systems (Hall *et al.*, 2011). Fluctuating selection cannot always proceed, though. When novel phenotypes correspond to increased generalism we do not expect past phenotypes to recur (Agrawal and Lively, 2002; Gandon *et al.*, 2008) since they will no longer be adaptive. Such expanding generalism during coevolution has been seen in other host-phage systems (Buckling and Rainey, 2002). Thus the relevance of fluctuating selection depends on the nature of the host-phage immune-infective phenotype interaction.

Another possible driver of coexistence are costs incurred by tradeoffs between growth and immunity (for host) or host range and immune evasion (for phage) (Chao *et al.*, 1977; Jover *et al.*, 2013; Levin *et al.*, 1977; Meyer *et al.*, 2016). A tradeoff between immunity and growth rate in the host can lead to the maintenance of a susceptible host population on which phages can persist (Chao *et al.*, 1977; Jones and Ellner, 2007; Jover *et al.*, 2013; Lenski and Levin, 1985; Levin and Udovic, 1977; Yoshida *et al.*, 2007). Tradeoffs often imply a high cost of immunity that does not always exist (e.g. Schrag and Mittler, 1996), particularly in the case of intracellular host immunity, as we show later.

Finally, in large host populations typical of bacteria, even low rates of immune loss could produce a substantial susceptible host subpopulation, which, in turn, could support phage reproduction and coexistence. Such loss of function in the host defenses could be due to either mutation or stochastic phenotypic changes. Delbrück (1946) initially described this hypothesis of loss of defense via back-mutation in order to challenge the evidence for lysogeny. Lenski (1988) reiterated this hypothesis in terms of phenotypic plasticity and noted that conditioning the production of a susceptible host population on a resistant one could lead to very robust, host-dominated coexistence. More recently, Meyer *et al.* (2012) presented an empirical example of a system in which stochastic phenotypic loss of resistance leads to persistence of a coevolving phage population.

We hypothesize that coexistence equilibria will be more robust under an immune loss mechanism than under a tradeoff mechanism (Lenski, 1988). We build a general mathematical model to demonstrate this point and then use a combination of experimental evidence and simulation-based modeling to apply this result to the coevolution of *Streptococcus thermophilus* and its lytic phage 2972 in the context of CRISPR immunity.

## 2. General Immune Loss Model

We begin with a general model that considers two populations of host (“defended” with a functional immune system; “undefended” without) and one population of pathogen. Starting from classical models of bacteria-phage dynamics (Levin *et al.*, 1977; Weitz, 2016), we add key terms to capture the effects autoimmunity (i.e., a tradeoff), immune loss, and the implicit effects of coevolution. This relatively simple model allows us to analyze steady states and parameter interactions analytically. Later, we examine the CRISPR-Cas immune system in detail and build a model with explicit coevolutionary dynamics.

We examine the chemostat system with resources:

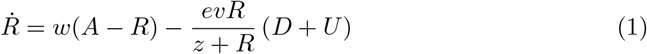

defended host:

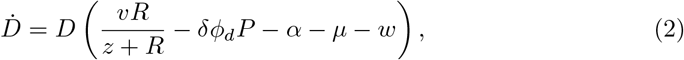

undefended host:

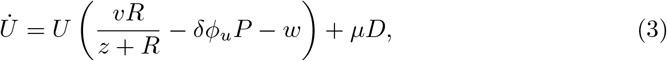

and phage:

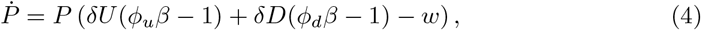

where parameter definitions and values can be found in Table 1 and rationale/references for parameter values in Supplementary Text S2. However, we describe here the parameters of direct relevance to coexistence.

**Table 1:** Definitions and oft used values/initial values of variables, functions, and parameters for the general mathematical model

| Symbol | Definition | Value |
| --- | --- | --- |
| $R$ | Resources | $R(0)=350 \quad \mu \text{ g/mL}$ |
| $D$ | Defended Host | $D(0)=10^6 \text{ cells/mL}$ |
| $U$ | Undefended Host | $U(0)=10^2 \text{ cells/mL}$ |
| $P$ | Phage | $P(0)=10^6 \text{ particles/mL}$ |
| $e$ | Resource consumption rate of growing bacteria | $5 \times 10^{-7} \quad \mu \text{ g/cell}$ |
| $v$ | Maximum bacterial growth rate | 1.4 divisions/hr |
| $z$ | Resource concentration for half-maximal growth | 1 $\mu \text{ g/mL}$ |
| $A$ | Resource pool concentration | 350 $\mu \text{ g/mL}$ |
| $w$ | Flow rate | 0.3 mL/hr |
| $\delta$ | Adsorption rate | $10^{-8} \text{ mL per cell per phage per hr}$ |
| $\beta$ | Burst Size | 80 particles per infected cell |
| $\phi_u$ | Degree of susceptibility of undefended host | 1 |
| $\phi_d$ | Degree of susceptibility of defended host | 0 |
| $\alpha$ | Autoimmunity rate | $2.5 \times 10^{-5} \text{ deaths per individual per hr}$ |
| $\mu$ | Rate of immune inactivation/loss | $5 \times 10^{-4} \text{ losses per individual per hr}$ |

First, we allow for defended host to come with the tradeoff of autoimmunity (*α*), which applies naturally to the CRISPR-Cas system examined later. While autoimmunity could either decrease the host growth rate (Vercoe *et al.*, 2013) or be lethal, we focus on the latter as lethality will increase the stabilizing effect of this tradeoff (Dy *et al.*, 2013; Paez-Espino *et al.*, 2013; Vercoe *et al.*, 2013). However, we also find similar general results when applying a penalty to the resource affinity or maximum growth rate of the defended host (Supplementary Text S1, Supplementary Figs 1-8).

Second, we add flow from the defended to undefended host populations representing loss of immunity at rate *µ*.

Finally, we model the effect of coevolution by allowing a fraction of even the defended host population to remain susceptible (0 *< ϕ*_*d*_ ≤ 1). In a symmetric fashion, even nominally undefended host may have secondary defenses against phage (0 *< ϕ*_*u*_ ≤ 1).

We analyze our model analytically as well as numerically to verify which equilibria are reachable from plausible (e.g., experimental) starting values (Supplementary Text S3).

Assuming no phage coevolution (*ϕd* = 0), this model has a single analytic equilibrium in which all populations coexist (Supplementary Table 1). In Fig 1, we explore model behavior under varying rates of autoimmunity (*α*) and immune loss (*µ*). Clearly when autoimmunity and loss rates surpass unity, defended host go extinct in the face of excessive immune loss and autoimmune targeting. At the opposite parameter extreme, we see coexistence disappear from the numeric solutions (Fig 1b) as phage populations collapse. This leads to a band of parameter space where coexistence is possible, stable, and robust. In this band, autoimmunity and/or immune loss occur at high enough rates to ensure maintenance of coexistence, but not so high as to place an excessive cost on immunity. Crucially, this band is much more constrained in the *α*-dimension, with autoimmunity restricted to an implausibly high and narrow region of parameter space. This suggests a greater robustness of coexistence under an immune loss mechanism even at low loss rates (Fig 1, Supplementary Figs 2-8). To assess more directly the degree of robustness of each driver of coexistence we can perturb our system and see its response. We move our system away from equilibrium 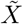 so that 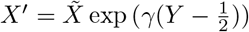 where *Y* ∼ Uniform[0, 1], and then solve numerically using *X*^′^ as our initial condition. Under increasing levels of perturbation the system is less likely to reach stable coexistence, specifically in the *α*-dimension, indicating that autoimmunity produces a far less robust coexistence regime (Fig 1c-e, Supplementary Figs 2-8).

**Figure 1:**
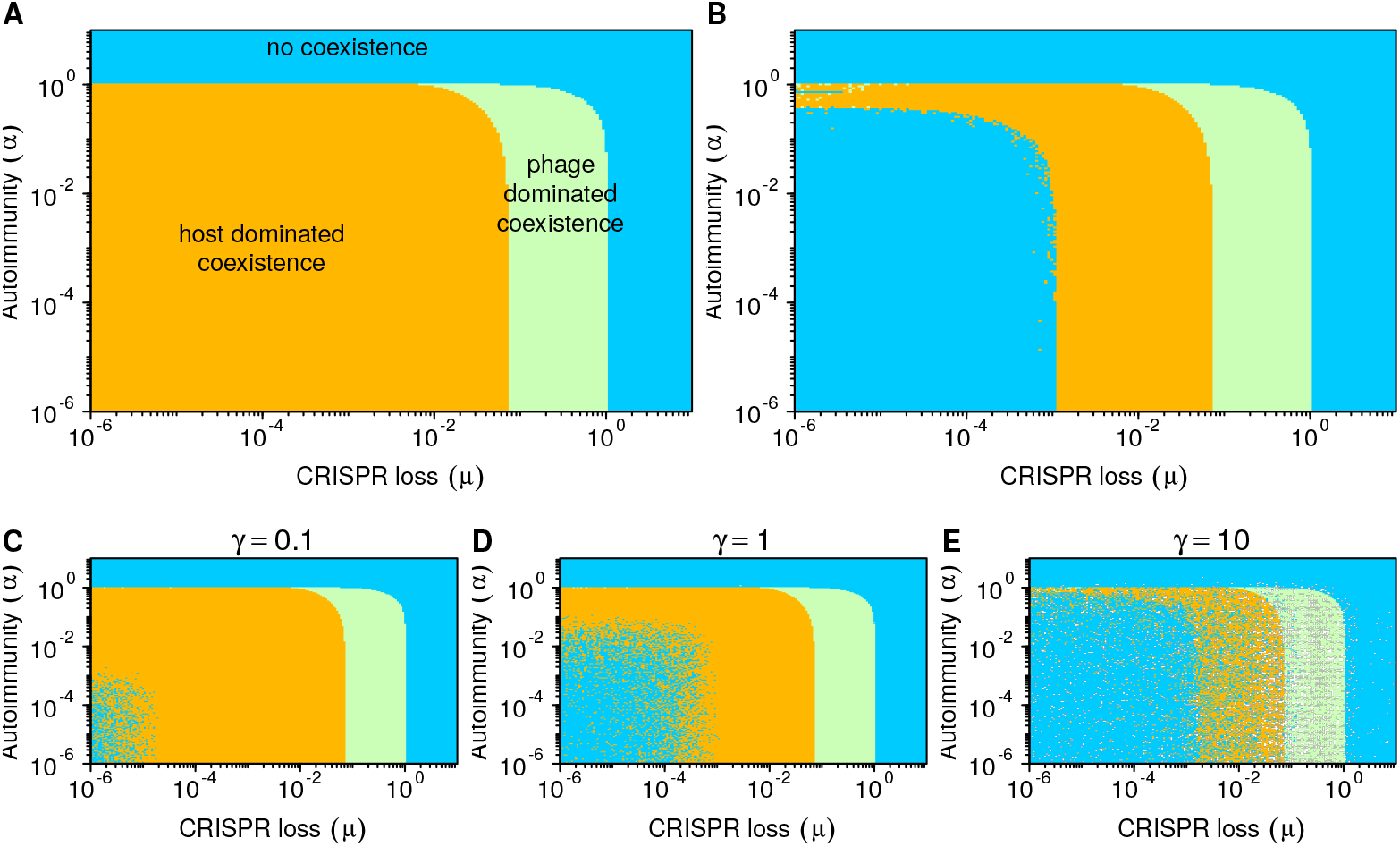
Model behavior under variations in the rates of autoimmunity (*α*) and CRISPR-Cas system loss. (*µ*) Equilibria (Supplementary Table 1) derived from Equations 1-4 are shown in (a) where orange indicates a stable equilibrium with all populations coexisting and defended host dominating phage populations, green indicates that all populations coexist but phages dominate, and blue indicates that defended bacteria have gone extinct but phages and undefended bacteria coexist. In (b) we find numerical solutions to the model at 80 days using realistic initial conditions more specific to the experimental setup (*R*(0) = 350, *D*(0) = 10^6^, *U* (0) = 100, *P* (0) = 10^6^). In this case orange indicates coexistence at 80 days with defended host at higher density than phages, green indicates a phage-dominated coexistence at 80 days, and blue indicates that coexistence did not occur. Numerical error is apparent as noise near the orange-blue boundary. We neglect coevolution and innate immunity in this analysis (*ϕu* = 1, *ϕd* = 0). (c-e) Phase diagrams with perturbed starting conditions. Numerical simulations with starting conditions (*X*(0) = [*R*(0)*, D*(0)*, U*(0)*, P* (0)]) perturbed by a proportion of the equilibrium condition 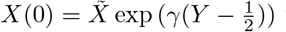 where *Y* ∼ *U*[0, 1] and *X*^˜^ signifies an equilibrium value to explore how robust the equilibria are to starting conditions. A single simulation was run for each parameter combination.

If we add large amounts of innate immunity to undefended host (*ϕu <* 0.5), we find phage-dominated coexistence for a wider range of *α* (Supplementary Fig 10). This result is in line with the counterintuitive suggestion that higher immunity may increase phage density by allowing the host population to increase in size (Iranzo *et al.*, 2013). However, secondary defense has minimal effects for more plausible levels of protection (*ϕu* closer to 1).

In the case of phage coevolution (*ϕd >* 0), the equilibria still have closed forms, but are not easily representable as simple equations and so are not written here. When 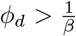, defended host contribute positively to phage growth, eventually shifting the coexistence equilibrium from host to phage dominance (Supplementary Fig 9).

## 3. A Case Study: CRISPR-Phage Coevolution

The CRISPR (Clustered Regularly Inter-spaced Short Palindromic Repeats) prokaryotic adaptive immune system incorporates specific immune memory in the form of short sequences of DNA acquired from foreign genetic elements (“spacers”) and then uses this memory to target the corresponding sequences (“protospacers”) during subsequent infections (Barrangou *et al.*, 2007; Bolotin *et al.*, 2005; Garneau *et al.*, 2010; Mojica *et al.*, 2005). CRISPR can lead to rapidly escalating arms races between bacteria and phages (Deveau *et al.*, 2008; Horvath *et al.*, 2008; Paez-Espino *et al.*, 2015), in which evolutionary and population dynamics occur on the same timescale (Childs *et al.*, 2012; Desai *et al.*, 2007; Gerrish and Lenski, 1998; Paez-Espino *et al.*, 2015).

CRISPR-Cas can quickly drive phages extinct in an experimental setting (van Houte *et al.*, 2016), but in some cases long-term CRISPR-phage coexistence has been observed (Paez-Espino *et al.*, 2015). Previous theoretical and limited experimental work has explained short-term coexistence through tradeoffs and spacer loss (Bradde *et al.*, 2017), and long-term coexistence by invoking continued coevolution via fluctuating selection (Childs *et al.*, 2012) or tradeoffs with host switching to a constitutive defense strategy such as surface receptor modification (Chabas *et al.*, 2016; Westra *et al.*, 2015).

However, these previous hypotheses are insufficient to explain simple coevolution experiments with *Streptococcus thermophilus* (type II-A CRISPR-Cas system) and its lytic phage 2972 resulting in long-term coexistence (Paez-Espino *et al.*, 2013, 2015). In these experiments, bacteria are resource-limited and appear immune to phages, implying they have “won” the arms race and that phages are persisting on a small susceptible subpopulation of hosts. Deep sequencing of the same experimental system shows dominance by a few spacers that drift in frequency over time, inconsistent with a fluctuating selection dynamic (Paez-Espino *et al.*, 2013). Specifically, these results contradict the coexistence regime seen in the Childs et al. (2014; 2012) model, wherein host are phage-limited and the system undergoes a fluctuating selection dynamic. Thus either (1) costs associated with CRISPR immunity or (2) the loss of CRISPR immunity is playing a role in maintaining susceptible host subpopulations on which phages can persist.

In this system, the primary cost of a functional CRISPR-Cas system is autoimmunity via the acquisition of self-targeting spacers. It is unclear how or if bacteria distinguish self from non-self during the acquisition step of CRISPR immunity (Kumar *et al.*, 2015; Levy *et al.*, 2015; Stern *et al.*, 2010; Wei *et al.*, 2015; Yosef *et al.*, 2012). In *S. thermophilus*, experimental evidence suggests that there is no mechanism of self vs. non-self recognition and that self-targeting spacers are acquired frequently (Wei *et al.*, 2015), which implies that autoimmunity may be a significant cost.

Outright loss of CRISPR immunity at a high rate could also lead to coexistence. The bacterium *Staphylococcus epidermidis* loses phenotypic functionality in its CRISPR-Cas system, either due to wholesale deletion of the relevant loci or mutation of essential sequences (i.e. the leader sequence or *cas* genes), at a rate of 10^-4^ -10^-3^ inactivation/loss events per individual per generation (Jiang *et al.*, 2013). Functional CRISPR loss has been observed in other systems as well (Garrett *et al.*, 2011; Palmer and Gilmore, 2010).

Below we replicate the serial-transfer coevolution experiments performed by Paez-Espino et al. (2013; 2015) and develop a simulation-based coevolutionary model to explain the phenomenon of coexistence.

### 3.1 Experiments

We performed long-term daily serial transfer experiments with *S. thermophilus* and its lytic phage 2972 in milk, a model system for studying CRISPR evolution (see Supplementary Text S4 for detailed methods). We measured bacteria and phage densities on a daily basis. Further, on selected days we PCR-amplified and sequenced the CRISPR1 and CRISPR3 loci, the two adaptive CRISPR loci in this bacterial strain.

From the perspective of density, phages transiently dominated the system early on, but the bacteria quickly took over and by day five appeared to be resource-limited rather than phage-limited (Fig 2a,b). This switch to host-dominance corresponded to a drop in phage populations to a titer two to three orders of magnitude below that of the bacteria. Once arriving at this host-dominated state, the system either maintained quasi-stable coexistence on an extended timescale (over a month and a half), or phages continued to decline and went extinct relatively quickly (Fig 2a,b). We performed six additional replicate experiments which confirmed this dichotomy between either extended coexistence (4 lines quasi-stable for *>* 2 weeks) or quick phage extinction (2 lines *<* 1 week) (Supplementary Fig 11).

Sequencing of the CRISPR1 and CRISPR3 loci revealed the rapid gain of a single spacer (albeit different spacers in different sequenced clones) in CRISPR1 followed by minor variation in spacer counts with time (Supplementary Fig 12), with CRISPR1 being more active than CRISPR3. We tracked the identity of the first novel spacer in the CRISPR1 array over time. We found a cohort of four spacers that persisted over time and were repeatedly seen despite a small number of samples taken at each time point (less than 10 per time point; Table 2). Other spacers were sampled as well, but this small cohort consistently reappeared while other spacers were only found at one or two timepoints, indicating this cohort was dominating the system (Supplementary Table 2). Such a pattern is inconsistent with a fluctuating selection hypothesis. Further, we did not observe frequent spacer loss in the CRISPR1 or CRISPR3 arrays.

**Table 2:** Sequencing data shows four first-order spacers that persist as a high-frequency cohort over time. Samples identified by the first novel spacer added to the array as compared to the wild-type. See Table S2 for complete spacer dynamics.

|  | Spacer ID |  |  |  |  |  |  |
| --- | --- | --- | --- | --- | --- | --- | --- |
| Time | D | E | F | G | Total in Cohort | Total Sampled Sequences | Percent Samples in Cohort |
| 1 | 0 | 0 | 0 | 0 | 0 | 3 | 0 |
| 2 | 1 | 1 | 1 | 0 | 3 | 4 | 75 |
| 3 | 2 | 0 | 1 | 1 | 4 | 5 | 80 |
| 4 | 0 | 0 | 0 | 0 | 0 | 1 | 0 |
| 5 | 0 | 0 | 2 | 1 | 3 | 7 | 43 |
| 11 | 1 | 2 | 0 | 2 | 5 | 7 | 71 |
| 15 | 1 | 1 | 1 | 0 | 3 | 7 | 43 |
| 25 | 0 | 5 | 0 | 0 | 5 | 8 | 63 |
| 35 | 0 | 1 | 0 | 2 | 3 | 6 | 50 |
| 40 | 0 | 0 | 0 | 2 | 2 | 9 | 22 |

### 3.2 CRISPR-phage Coevolutionary Model

We next built a hybrid deterministic/stochastic lineage-based model similar to an earlier model by Childs et al. (2014; 2012) that explicitly models the coevolutionary dynamics of the CRISPR-phage system wherein bacteria acquire spacers to gain immunity and phages escape spacers via mutations. Our simulations also replicate the resource dynamics of a serial dilution experiment, wherein the system undergoes large daily perturbations.

We model phage mutations only in the protospacer adjacent motif (PAM) region, which is the dominant location of CRISPR escape mutations (Paez-Espino *et al.*, 2015) to prevent the possibility of spacer re-acquisition. This approach differs from previous models which considered mutations in the protospacer region itself (e.g. Childs *et al.*, 2012; Iranzo *et al.*, 2013; Weinberger *et al.*, 2012) and thus allowed for the possibility of spacer re-acquisition. We justify modeling only PAM mutations with three arguments. First, the probability of spacer re-acquisition will be quite low if there are many protospacers. Second, re-acquired spacers will already have undergone selection for escape mutation by phage, and, assuming that there are therefore diverse escape mutations in the phage population, these spacers will thus provide limited benefit to the host. Third, as we move away from the PAM along the protospacer sequence, more substitutions are tolerated by the CRISPR matching machinery (Semenova *et al.*, 2011), meaning that mutations farther away from the PAM will be less effective at escaping immunity (Martel and Moineau, 2014).

We model population dynamics using differential equations for resources:

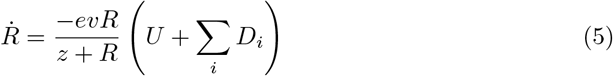

CRISPR-enabled bacteria with spacer set *X*_*i*_

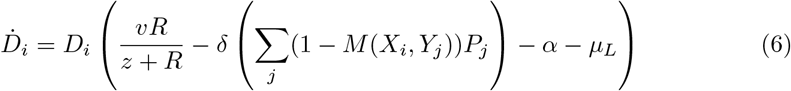

a pool of undefended bacteria with a missing or defective CRISPR-Cas system:

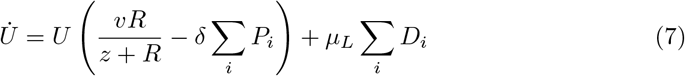

and phages with protospacer set *Yi*:

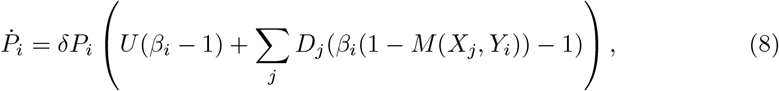

and stochastic events occur according to a Poisson process with rate λ

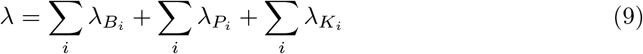

which is a sum of the total per-strain spacer-acquisition rates:

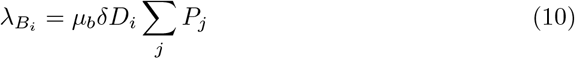

total per-strain PAM mutation rates:

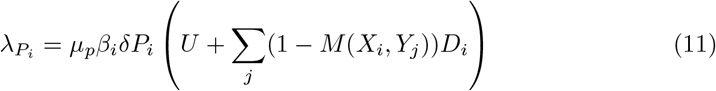

and total per-strain PAM back mutation rates:

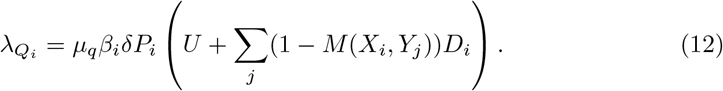

In this way each unique CRISPR genotype (*Xi*), defined as a set of linked spacers sharing the same array, is modeled individually, as is each phage genotype (*Yi*). As new spacers are added and new PAMs undergo mutation, new pairs of genotypes and equations are added to the system. Host that have undergone immune loss are modeled separately (*U*), as if they have no CRISPR-Cas system.

The function *M*(*Xi, Yj*) is a binary matching function between (proto)spacer content of bacterial and phage genomes that determines the presence or absence of immunity. We refer to the “order” of a host or phage strain, which is the number of evolutionary events that strain has undergone, |*X*_*i*_ | or *n*_*s*_ - |*Y*_*i*_ | respectively. The PAM back mutation rate *µ*_*q*_ describes the rate at which we expect a mutated PAM to revert to its original sequence (assuming the mutation is a substitution). While back mutation is not required to generate stable host-dominated coexistence, it greatly expands the relevant region of parameter space because it allows phages to avoid the cost we will impose on PAM mutations, discussed below, when those immune escape mutations are no longer beneficial. Recombination among viral strains could have a similar effect by providing another route to an un-mutated or less mutated genome. Páez-Espino (2015) suggest that recombination can produce stable host-dominated coexistence, although we reject such diversity-driven hypotheses (e.g. Childs *et al.*, 2012) based on our sequencing data.

We assume that the number of PAM mutations in a single phage genome is constrained by a tradeoff with phage fitness, as this is necessary to prevent the total clearance of protospacers from a single strain at high mutation rates. Increases in host breadth at the species level generally come at a cost for viruses due to pleiotropic effects (Ferris *et al.*, 2007). More broadly, mutations tend to be deleterious on average (e.g. Chao, 1990). It is reasonable to speculate that phages have evolved under pressure to lose any active PAMs on their genomes, and thus that the persisting PAMs may have been preserved because their loss is associated with a fitness cost.

The function

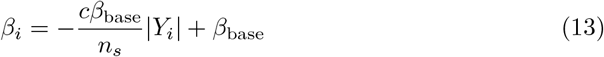

incorporates a linear cost of mutation into the phage burst size. See Table 3 for further definitions of variables, functions, and parameters in Equations 5-13. Simulation procedures and rationale for parameter values, including phage genome size, are detailed in Supplementary Text S3.

**Table 3:**
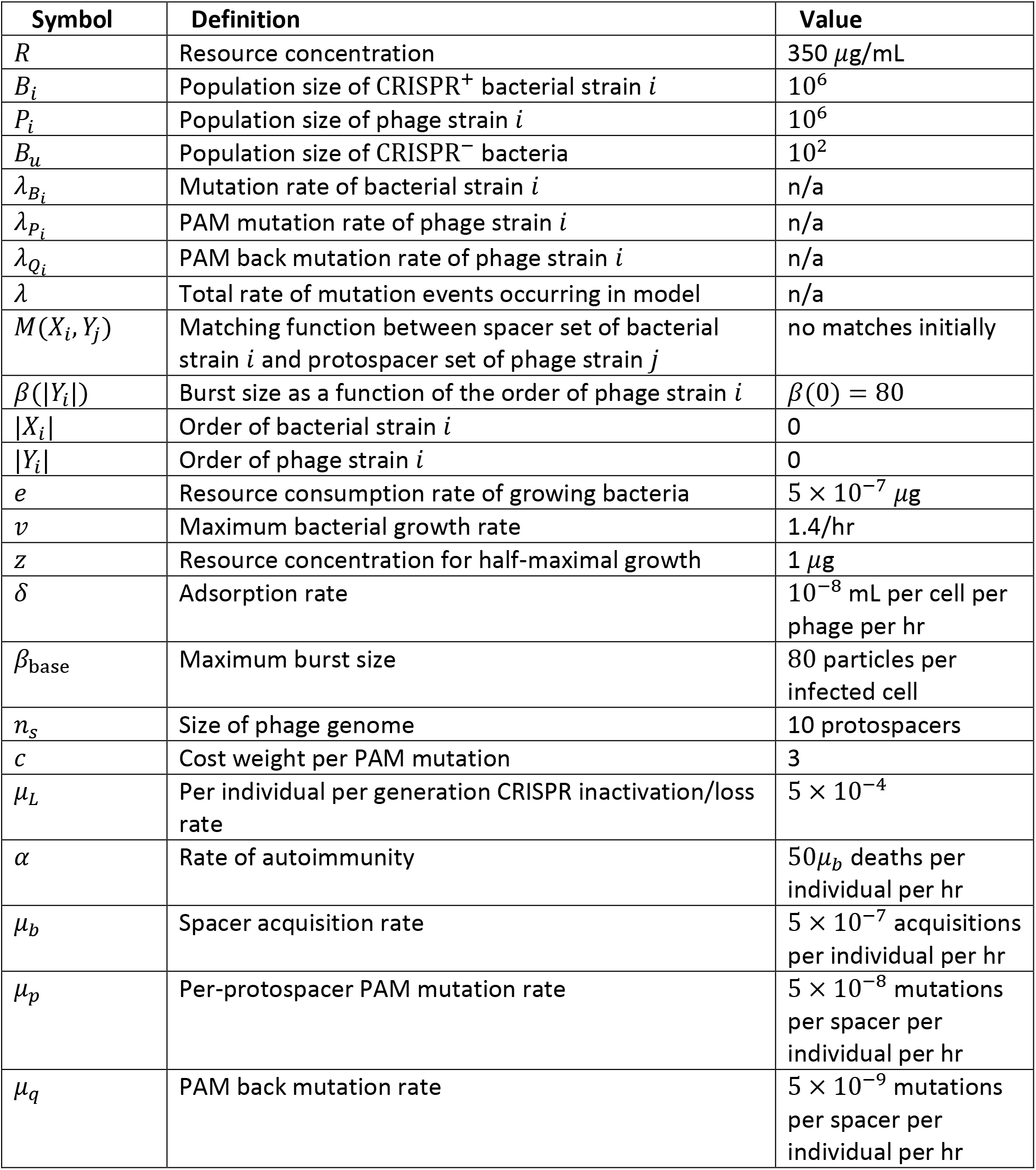
Definitions and oft used values/initial values of variables, functions, and parameters for the simulation model

#### 3.2.1 Stable Host-Dominated Coexistence

Simulations with immune loss reliably produce extended coexistence within a realistic region of the parameter space (Fig 3) thus replicating our experimental results (Fig 2), and confirming our qualitative results from the simpler deterministic model (Fig 1). We observed no simulations in which autoimmunity alone produced stable coexistence. This agrees with our earlier numerical results from the general model where unrealistically high rates of autoimmunity were required to produce coexistence.

**Figure 2:**
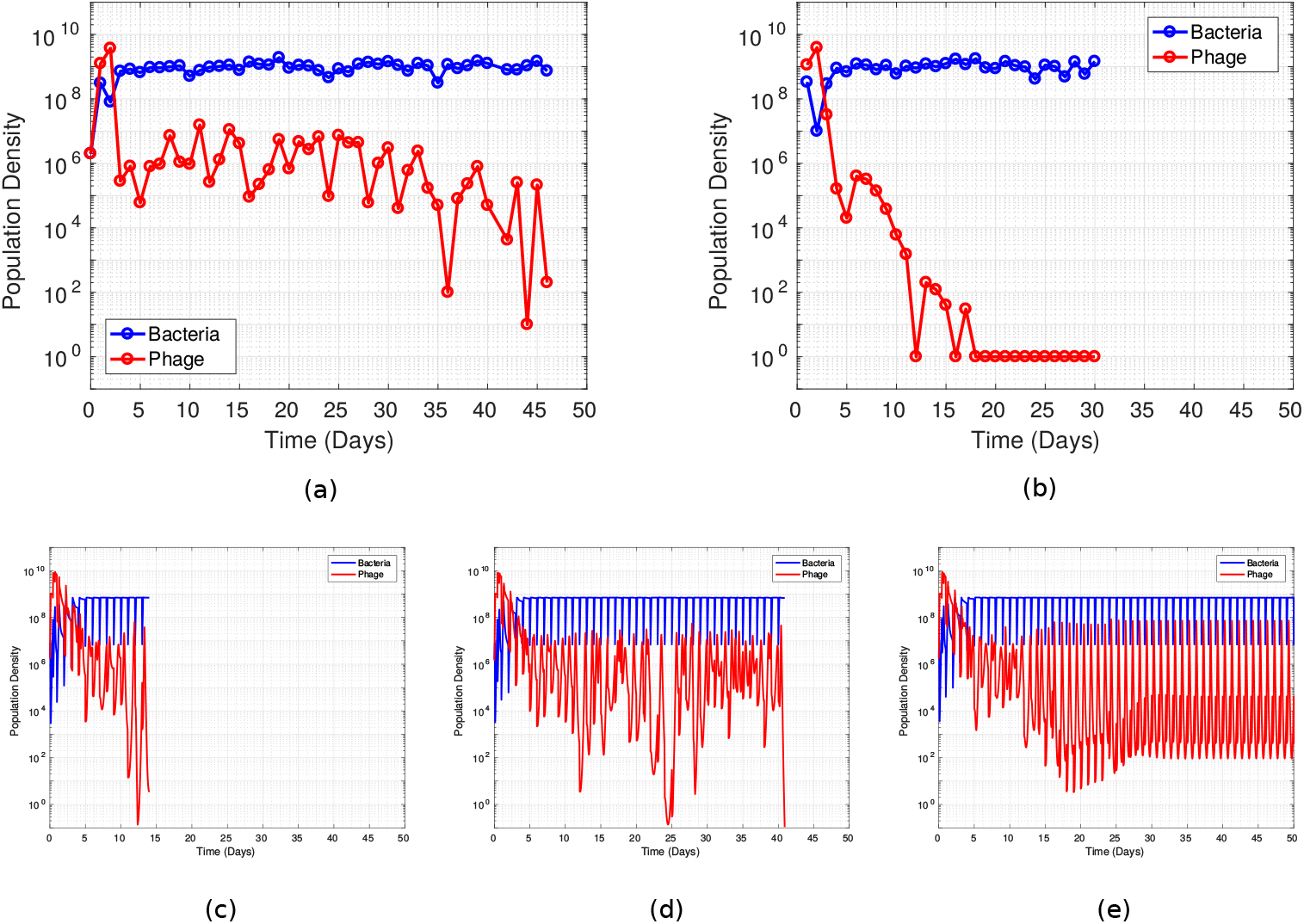
Serial transfer experiments carried out with *S. thermophilus* and lytic phage 2972. Bacteria are resource-limited rather than phage-limited by day five and phages can either (a) persist at relatively low density in the system on long timescales (greater than 1 month) or (b) collapse relatively quickly. These results agree with those of Paez-Espino (2015) where coexistence was observed in *S. thermophilus* and phage 2972 serially transferred culture for as long as a year. Experiments were initiated with identical starting populations and carried out following the same procedure. In (c-e) we show that our simulations replicate the qualitative patterns seen in the data, with an early phage peak, followed by host-dominated coexistence that can either be (c) stable, (d) sustained but unstable, or (e) short-lived. Each plot is a single representative simulation and simulations were ended when phages went extinct. Note that experimental data has a resolution of one time point per day, preventing conclusions about the underlying population dynamics (e.g., cycling), whereas simulations are continuous in time.

**Figure 3:**
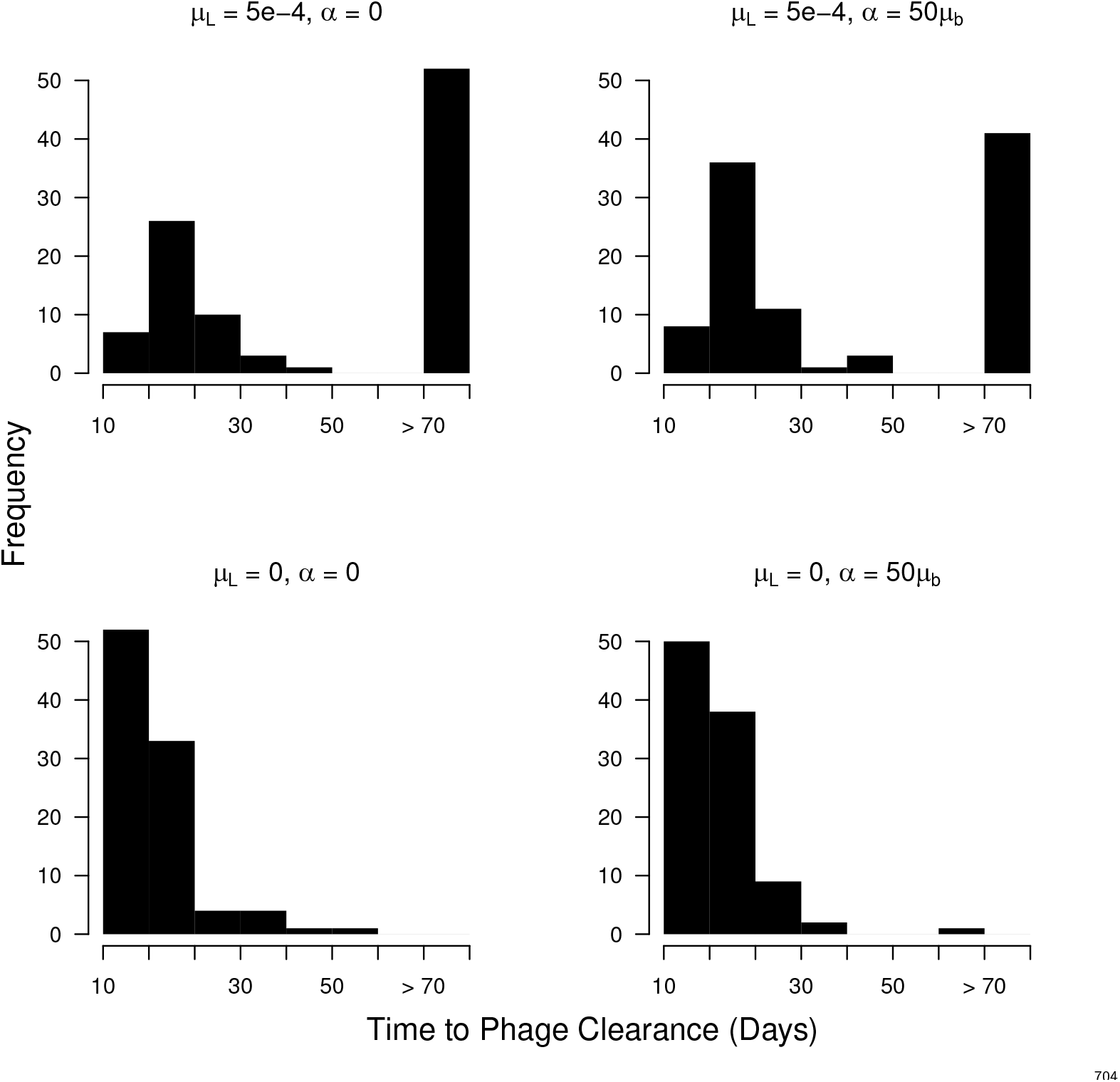
Distribution of phage extinction times in bacterial-dominated cultures with different possible combinations of coexistence mechanisms. The peak at ≥ 75 corresponds to what we call stable coexistence (simulations ran for a maximum of 80 days). There is no significant difference between the top two panels in the number of simulations reaching the 80 day mark (*χ*^2^ = 2.8904, *df* = 1, *p* - value = 0.08911). Back mutation was set at *µq* = 5 × 10^-9^.

Similar to our experimental results, for a single set of parameters this model can stochastically fall into either stable coexistence or a phage-free state (Fig 3). The relative frequencies with which we see each outcome, as well as the distribution of times that phages are able to persist, depend on the specific set of parameters chosen. In particular, increasing the PAM back mutation rate will increase the probability of the coexistence outcome (Fig 4), although even in the absence of back mutation the system will occasionally achieve stable coexistence. This dependence on back mutation is caused by the combined effects of the cumulative cost we impose on PAM mutations and the inability of phages to keep up with host in a continuing arms race. In the early stages of the arms race it is optimal for phages to continue undergoing PAM mutations as the most abundant available hosts are high-order CRISPR variants, whereas once hosts are able to pull sufficiently ahead of phages in the arms race it becomes optimal for phages to feed on the lower-density but consistently available CRISPR-lacking host population (Supplementary Fig 13).

**Figure 4:**
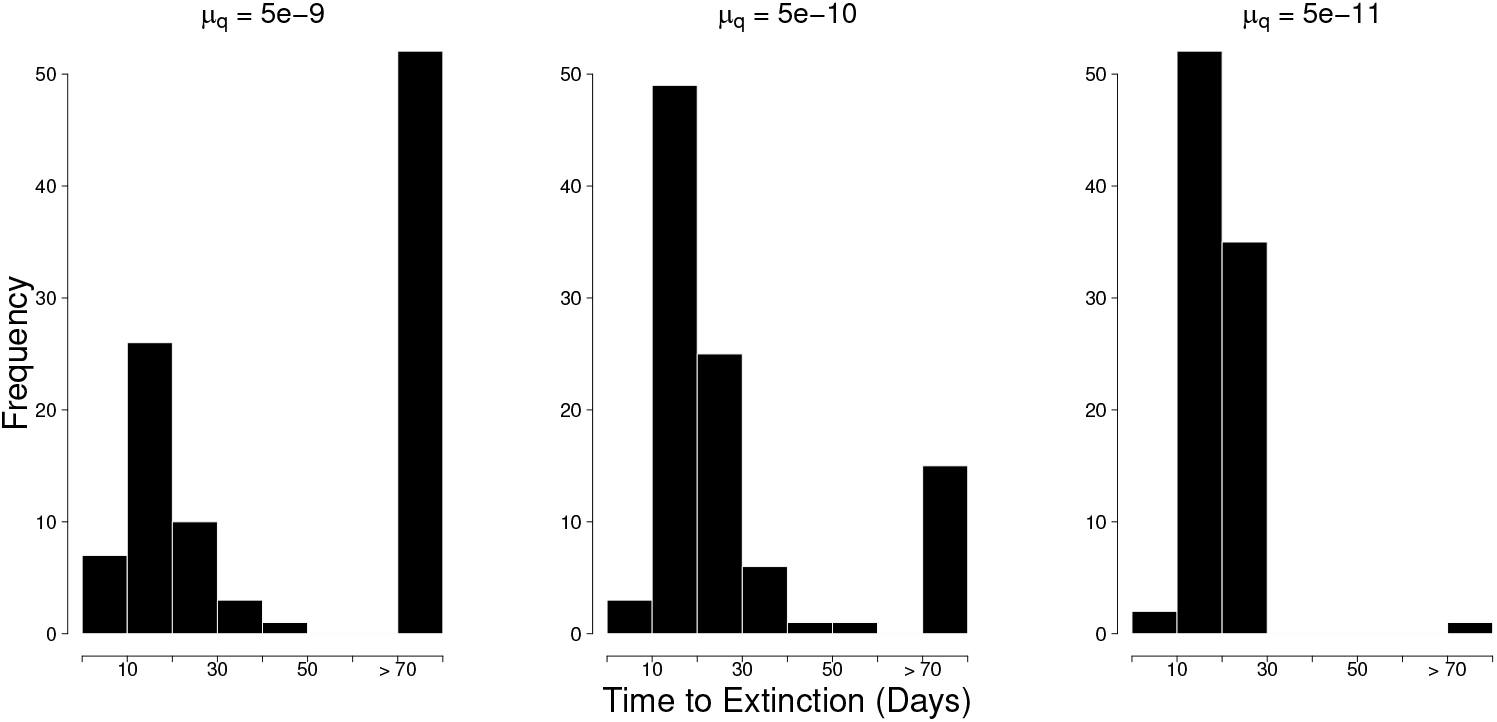
Distribution of phage extinction times in bacterial-dominated cultures with different rates of PAM back mutation in phages. (*µ*_*q*_) The peak at 80 corresponds to what we call stable coexistence (simulations ran for a maximum of 80 days). These results are shown for a locus-loss mechanism only (*µL* = 5 × 10^-4^, *α* = 0). The histogram for *µ q* = 5 × 10 ^- 8^ is omitted as it is nearly identical to that for *µq* = 5 × 10^-9^, indicating that the height of the coexistence peak saturates at high back mutation.

The adsorption rate, on a coarse scale, has an important effect on how the model behaves (Supplementary Fig 14). At high values of *δ* where we would expect phages to cause host extinction in the absence of CRISPR immunity (*δ* = 10^-7^) we see that long-term coexistence occurs rarely, and is negatively associated with the phage back mutation rate. In this case phages will rapidly consume the susceptible host population and crash to extinction unless they have undergone PAM mutations that lower their growth rate. This causes a reversal in the previous trend seen with back mutation where the ability of phages to escape the costs of PAM mutation was essential to their persistence. A decrease in the adsorption rate to a very low value (*δ* = 10^-9^) leads to most simulations persisting in host-dominated coexistence until the 80 day cutoff. Because both evolutionary and demographic dynamics occur much more slowly in this case, long term persistence does not necessarily imply actual stability, as suggested by our and previous (Paez-Espino *et al.*, 2015) experimental results in which coexistence eventually ends. In general, lower adsorption rates lead to longer periods of host-dominated coexistence and reduce the chance of phage extinction.

The failure of autoimmunity to produce coexistence warrants further investigation. Upon closer examination, it is clear that in the early stages of the arms race where CRISPR-enabled bacteria have not yet obtained spacers or been selected for in the host population, phages are able to proliferate to extremely high levels and greatly suppress the CRISPR-lacking host. Because autoimmunity as a mechanism of coexistence relies on the continued presence of immune-lacking host, it may not be able to function in the face of this early phage burst if susceptible host are driven extinct. There is a possibility that very low locus loss rates that reintroduce CRISPR-lacking bacteria but do not appreciably contribute to their density combined with high rates of autoimmunity could maintain high enough density susceptible host populations to sustain phage. To investigate this possibility we imposed a floor of *U >* 1 and ran further simulations. Even with very high rates of autoimmunity based on an upper limit of likely spacer acquisition rates (*α* = 50*µb*, *µb* = 10^-5^) the susceptible host population does not grow quickly enough to sufficiently high levels to sustain phage (Supplementary Fig 15). Thus it is not early dynamics that rule out autoimmunity but the insufficiency of the mechanism itself for maintaining large enough susceptible host populations.

#### 3.2.2 Transient Coexistence with Low Density Phage

While we do not observe stable coexistence in any case where there is not loss of the CRISPR-Cas immune system, we did observe prolonged phage persistence in some cases where *µL* = *α* = 0 (Fig 3) and in cases with autoimmunity only (*µL* = 0). Phages were able to persist at very low density (∼ 10 - 100 particles/mL) for as long as two months in a host-dominated setting without the presence of a CRISPR-lacking host subpopulation (Fig 3, Supplementary Fig 16). It appears that in these cases phages are at sufficiently low density as to have a minimal effect on their host population and thus that host strain is selected against very slowly. Because the phages have undergone many PAM mutations at this point they are unable to proliferate rapidly enough between dilution events to have an easily measurable impact on the host population. Essentially, phages delay their collapse by consuming their host extremely slowly (Supplementary Fig 16). However, with an active locus loss mechanism (i.e., *µL >* 0), we did not see this sustained but unstable coexistence occur, likely because the undefended hosts would have driven the phage population to higher levels and increased selection on the susceptible CRISPR variants.

## Discussion

We paired a general model of immunity with a case study of the CRISPR immune system to characterize and contrast the potential drivers of long-term host-phage coexistence in well-mixed systems. We found that a tradeoff mechanism does not lead to a robust coexistence equilibirum in the case of intracellular host immunity. We also ruled out coevolutionary drivers of coexistence in the *S. thermophilus*-phage 2972 system based on a combination of our own sequencing data and previous work on the same system (Paez-Espino *et al.*, 2013). Since some mechanism(s) must be producing susceptible hosts on which phages can replicate, we are left with an immune loss hypothesis as the best remaining explanation for our empirical results. Our simulations showed that the addition of early coevolutionary dynamics alongside immune loss replicates key features of our experimental results, including stochastic switching between the possible outcomes of long term coexistence and rapid phage clearance. Therefore we predict that that the CRISPR-Cas immune system is lost at a nontrivial rate in *S. thermophilus* in addition to *S. epidermidis* (Jiang *et al.*, 2013), and possibly other species.

With regards to CRISPR, while our experiments do not speak to the relative importance of locus loss versus costly autoimmunity, our theoretical results reject autoimmunity as a realistic mechanism of phage persistence. Our experimental setup was in serial dilution, which subjects the culture to large daily perturbations, ruling out any mechanism that does not produce a robust coexistence regime.

We emphasize that CRISPR immunity, and immunity in general, is still likely costly (Vale *et al.*, 2015). Nevertheless, in cases of intracellular host immunity those costs are insufficient to drive continued phage persistence in the environment. Intracellular immunity destroys phages rather than simply preventing phage replication. Thus the threshold density of susceptible host for phage persistence is higher than in systems where hosts have an extracellular defense strategy (i.e. receptor/envelope modification), meaning the cost of immunity must be higher. When hosts escape phage predation via receptor modifications, a growth-resistance tradeoff may lead to coexistence.

Our sequencing results in the *S. thermophilus* system reject coevolutionary mechanisms for coexistence. We can directly reject an arms race dynamic since it predicts the rapid, continued accumulation of spacers, which does not occur in our data. A fluctuating selection dynamic makes the more subtle prediction that the frequencies of spacers in the population should cycle over time, with different spacers dominating at different times. Even with relatively small sample sizes (<10 CRISPR loci sequenced per timepoint), we see a small cohort of spacers increase in frequency early in the experiment and continue to be detected at later timepoints (Table 2). These results are consistent with those of Paez-Espino et al. (2013) who performed deep sequencing with the same phage-host system and observed dominant spacers that drifted in frequency over time. This continued detection and dominance of particular spacers rules out strong fitness differences between these spacers, which, in turn, contradicts the expectation of fluctuating selection that fitnesses change over time. Our stochastic simulations agree, with coevolutionary dynamics in the absence of loss or cost most often yielding rapid phage extinction and only occasionally showing coexistence for over a month – but never exhibiting sustained coexistence (Fig 3).

A similar model by Childs et al. (2014) found that a fluctuating selection dynamic could lead to long term coexistence in a CRISPR-phage system when arrays were “saturated”, in the sense that they were filled to some preset maximum capacity with spacers, which we do not observe in our experimental data. The fact that we see little expansion of the array suggests that hosts are completely immune to phages, as rapid phage genome degradation inside the CRISPR-immune cells can prevent further uptake of spacers (Semenova *et al.*, 2016).

While we conclude that immune loss plays a key role in our system, it is not immediately clear why bacterial immune systems would lose functionality at such a high rate. Our sequencing of the *S. thermophilus* CRISPR loci did not reveal pervasive spacer loss events, indicating that immune loss is at the system rather than spacer level. Perhaps in the case of CRISPR there is some inherent instability of the locus, leading to high rates of horizontal transfer (Chakraborty *et al.*, 2010; Garrett *et al.*, 2011; Godde and Bickerton, 2006; Palmer and Gilmore, 2010; Shah and Garrett, 2011). Jiang *et al.* (2013) propose that CRISPR loss is a bet-hedging strategy that allows horizontal gene transfer to occur in stressful environments (e.g., under selection for antibiotic resistance). This proposal is consistent with evidence that CRISPR does not inhibit horizontal gene transfer on evolutionary timescales (Gophna *et al.*, 2015). A high rate of CRISPR loss and inactivation could produce pressure for bacteria to frequently acquire new CRISPR-Cas systems through horizontal gene transfer, perhaps explaining why strains with multiple CRISPR-Cas systems are frequently observed, including *S. thermophilus* (Cai *et al.*, 2013; Horvath *et al.*, 2009). This is consistent with a broader view in which prokaryotic defense systems appear to be labile, having higher rates of gain and loss than other genetic content (Puigbò *et al.*, 2017).

While some clear anecdotes of immune loss exist (Jiang *et al.*, 2013; Meyer *et al.*, 2012), other examples of this phenomenon may have been missed because it is difficult to detect. Phages will quickly destroy any evidence of loss, and loss rates can be low while still affecting population dynamics. Jiang et al. (2013) go to great lengths to demonstrate loss in their system. Particularly with complex systems like CRISPR-Cas, a mutation in any number of components can lead to inactivation, making loss hard to detect from genetic screens. Phenotypic screens like those of Jiang et al. (2013) require the engineering of CRISPR spacer content and/or plasmid sequence as well as an otherwise competent host.

Other paths to sustained coexistence between CRISPR-enabled hosts and phages may also exist. There is a great diversity of CRISPR-Cas system types and modes of action (Makarova *et al.*, 2015) and the particular mechanism of each system may lead to distinct host-phage dynamics. That being said, our model of CRISPR evolutionary dynamics is rather general, and we recovered similar qualitative results over a wide range of parameter values apart from the *S. thermophilus*-specific parameter space.

Finally, our results show that the regular loss of immunity can sustain a viable phage population, leading to the maintenance of selective pressure and thus keeping immunity prevalent in the population overall. Even though long-term coexistence with phages may not affect overall host population density, we suggest that, counterintuitively, the periodic loss of individual immunity may drive the maintenance of a high population immune prevalence.

## Acknowledgments

This work was supported by funding from the University of Maryland and the U.S. Department of Education GAANN program. This material is based upon work supported in part by the U. S. Army Research Laboratory and the U. S. Army Research Office under contract/grant number #W911NF-14-1-0490. SM acknowledges funding from the Natural Sciences and Engineering Research Council of Canada (Discovery program). SM holds a T1 Canada Research Chair in Bacteriophages. PLFJ was supported in part by NIH R00 GM104158.

